# Insights into Intestinal Barrier Disruption During Long-Term Gut *Chlamydia* Colonization in Mice: A Single-Cell Transcriptomic Approach

**DOI:** 10.1101/2024.12.02.626423

**Authors:** Yicun Jiang, Sheng Xie, Chuqiang Shu, Jiao Wan, Youyou Huang, Luying Wang, Qi Zhang, Zengzi Zhou, Xin Sun, Tianyuan Zhang, Qi Tian

## Abstract

*Chlamydia trachomatis*, an intracellular pathogen, stands as the most prevalent sexually transmitted bacterial infection among women globally. Traditionally recognized as a genital pathogen, recent research indicates that the gastrointestinal tract may also act as a reservoir for its long-term colonization. However, the mechanisms underlying *Chlamydia*’s ability to persist in the gut remain poorly understood. This gap in knowledge limits our ability to develop effective treatments for persistent *Chlamydia* infections. In this study we utilized single-cell RNA sequencing to analyze the gene expression profiles and cellular heterogeneity of mouse colonic tissues during *Chlamydia* long-term infection. This approach provided detailed insights into the transcriptional changes and cellular interactions involved in the persistence of *Chlamydia* in the gut. Our results revealed significant alterations in the gene expression profiles of various intestinal cell populations, with distinct molecular pathways contributing to *Chlamydia* persistence. Notably, we observed a reduction in the expression of markers associated with epithelial tight junctions, indicating a potential breakdown of the intestinal epithelial barrier. This impairment may facilitate the penetration of *Chlamydia* into deeper tissues and contribute to the initiation of infection. We also found dysregulation of the transcriptional networks in goblet cells and an imbalance in communication between immune and epithelial cells. These disruptions were linked to the pathogen’s ability to establish persistent colonization and infection.

**IMPORTANCE:** Few studies have explored *Chlamydia* persistence in the gastrointestinal tract. In this study, we use single-cell RNA sequencing to identify the molecular and cellular mechanisms driving the pathogen’s long-term colonization. Our findings provide crucial insights into how *Chlamydia* overcomes the host’s immune defenses and epithelial barriers to establish chronic infection in the gut. Notably, we identify disruption of epithelial tight junctions and an imbalance in immune-cell interactions, offering new avenues for therapeutic interventions aimed at restoring mucosal integrity and preventing persistent infection.

## INTRODUCTION

*Chlamydia trachomat* (*C. trachomatis*) is the most common sexually transmitted infection globally(1), with significant clinical consequences for women, including pelvic inflammatory disease and infertility(2). While traditionally considered a genital pathogen, *Chlamydia* has also been frequently detected in the gastrointestinal (GI) tract of various hosts, including humans(3, 4). Recent studies suggest that the gut may serve as a natural reservoir for long-term *Chlamydia* colonization(5, 6). Evidence shows that *Chlamydia* can spread from the genital tract to the large intestine, where it establishes persistent colonization(7). However, the impact of *Chlamydia* persistence in the GI tract remains underexplored. Some studies propose that persistent gut infection may act as a reservoir, facilitating *Chlamydia*’s spread and reinfection within the host(5). Some research suggests that co-infection of the gut and genital tract during the acute phase could trigger pathogenic immune responses in the upper genital tract(8, 9).

The GI tract, particularly the large intestine, plays a crucial role in absorption and feces formation. Pathogen infections in the gut may compromise the intestinal barrier, leading to immune dysregulation and contributing to a range of intestinal diseases(10, 11). As an obligate intracellular pathogen, *Chlamydia* has developed complex mechanisms to adhere to and invade host cells, including intestinal epithelial cells. Upon infection, *Chlamydia* stimulates cells to produce a broad spectrum of cytokines, thereby modulating host immune responses and inflammation(12). Within infected cells, *Chlamydia* exhibits diverse developmental forms, suggesting possible adaptive strategy to survive in different intestinal cell types(13). Despite these insights, the precise mechanisms by which *Chlamydia* colonizes and persists in the large intestine remain poorly understood. Limited research has focused on the molecular and cellular changes that occur within intestinal tissues during infection. Understanding how *Chlamydia* disrupts intestinal cell composition and the molecular basis of its persistence is critical for developing effective therapeutic strategies and identifying novel treatment targets.

To investigate these mechanisms, we employ single-cell RNA sequencing (scRNA-seq), a useful technique that allows for the detailed analysis of gene expression at the single-cell level(14). This technology reveals the cellular heterogeneity and dynamic changes within populations that traditional bulk sequencing methods cannot capture(15). By analyzing scRNA-seq data from *Chlamydia*-infected mouse colonic tissues, we aim to elucidate how *Chlamydia* infection alters the intestinal microenvironment and affects immune and epithelial cell populations.

This study centered around the hypothesis that *Chlamydia* infection induces changes in the intestinal cellular landscape, particularly within epithelial cells. These alterations may drive inflammation and tissue damage, further compromising gut health(16). Specifically, we aim to investigate the site of persistent *Chlamydia* infection in the GI tract and to characterize the changes in cell populations and gene expression profiles in infected GI tissues. We focused on classifying and analyzing epithelial cell populations, which play critical roles in mucosal immunity, and explore the transcriptional regulation networks within these cells that may underlie the pathogenic mechanisms of *Chlamydia* infection. Additionally, we will examine how disrupted cell-to-cell communication pathways may impair immune responses(17), and investigate the role of *Chlamydia* in modulating the differentiation and lineage progression of cells within the gut.

By investigating the cellular and molecular mechanisms that support *Chlamydia* persistence in the gut, our study contributes to a deeper understanding of gut-specific *Chlamydia* infections and may inform the development of future treatment strategies.

## RESULTS

### The large intestine serves as a primary site for long-term *Chlamydia* colonization in the gut

To investigate the dynamics of *Chlamydia muridarum* (*C.muridarum*)colonization in the GI tract, 20 mice were divided into five groups (four mice per group) and inoculated intragastrically with live *C. muridarum*. GI tissues were collected at five time points: day 3, day 7, day 14, day 28, and day 35 post-infection, with one group sacrificed at each time point. The GI tract was segmented into the stomach, small intestine (SI)—comprising the duodenum, jejunum, and ileum—and large intestine (LI), which includes the cecum, colon, and rectum. All gut contents were removed, and the tissues were homogenized to evaluate the infection burden.

At day 3 post-infection, *Chlamydia* was detected across all segments of the GI tract. Over the following weeks, the pathogen burden in the stomach and small intestine progressively declined, eventually clearing entirely on day 35, while the burden in the large intestine steadily increased. By day 35, *Chlamydia* was exclusively detected in the large intestine, indicating its role as the primary site for long-term colonization (Figure 1A).

**Figure 1:**
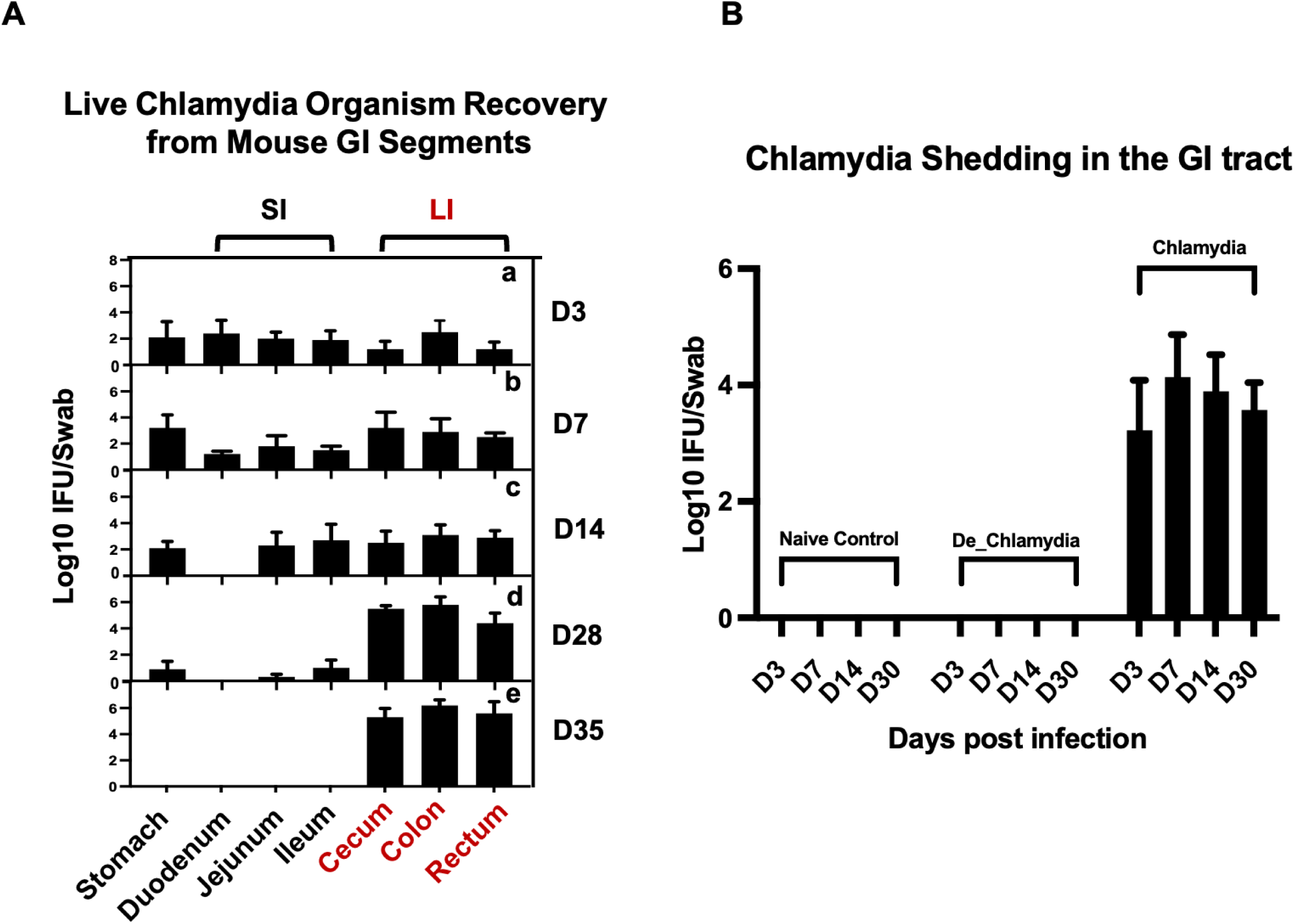
Localization of Long-Term *Chlamydia* Colonization in the Large Intestine of Mice. [A] Temporal progression of *Chlamydia* colonization across GI tract segments post-intragastric inoculation. Groups of C57BL/6J mice were euthanized on days 3 (a), 7 (b), 14 (c), 28 (d), and 35 (e) post-inoculation to assess the presence of live *Chlamydia* organisms within various tissue segments, including the stomach, small intestine (SI)—comprising duodenum, jejunum, ileum—and large intestine (LI), which includes the cecum, colon, and rectum. [B] Monitoring of *Chlamydia* shedding through analysis of rectal swabs collected from three different groups to evaluate the persistence of the pathogen in the GI tract over time.

To further study the impact of persistent *Chlamydia* infection on gut epithelial cells, three additional groups of mice (three per group) were inoculated with live *Chlamydia muridarum* (*Chlamydia*), heat-inactivated *Chlamydia* (De_*Chlamydia*), or SPG (Naive control). These groups were used for subsequent RNA sequencing analysis. The infection burden was monitored by recovering *Chlamydia* inclusions from rectal swabs. Persistent colonization was observed only in the group inoculated with live *Chlamydia*, with the pathogen detectable in rectal swabs up to day 30 post-infection. No *Chlamydia* was recovered from the rectal swabs of mice in the heat-inactivated *Chlamydia* or SPG control groups, confirming the necessity of viable *Chlamydia* for establishing persistent colonization (Figure 1B)

### Cellular Composition and Functional Changes in the Mouse Colon Following *Chlamydia* Infection

To explore the impact of *Chlamydia* infection on specific cell populations within the mouse colon, we conducted a comprehensive scRNA-seq analysis of colon tissues from mice infected with live *Chlamydia*, heat-inactivated *Chlamydia*, and uninfected controls. After initial data quality control, we obtained transcriptomic profiles from a total of 59,197 cells, which we normalized and subjected to principal component analysis and UMAP clustering. This analysis identified 27 distinct cell clusters (C0-26), which were annotated using known cellular markers (Figure 2A and S1A).

**Figure 2:**
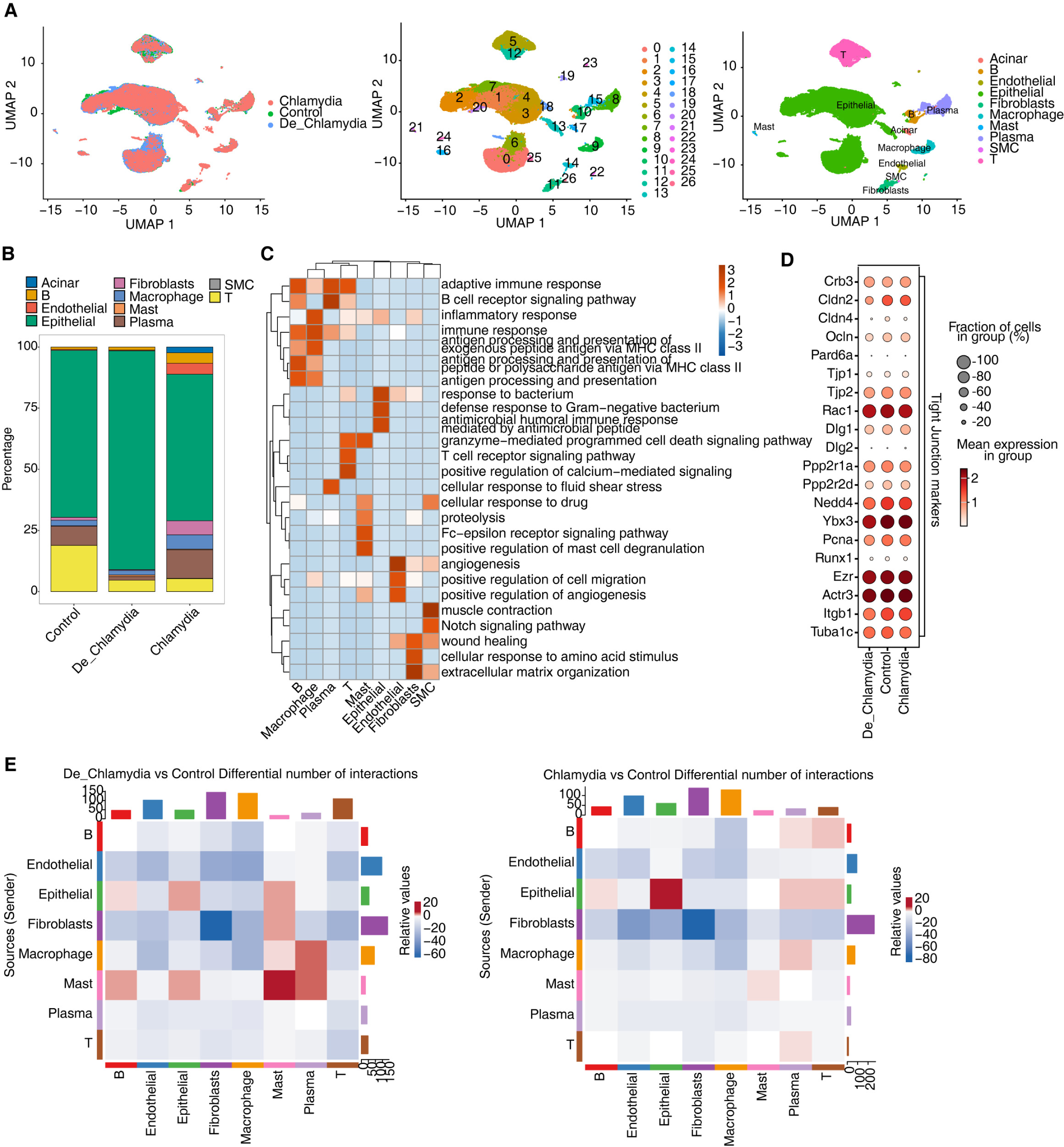
ScRNA-seq reveals *Chlamydia* infection associated shift in intestinal cell composition. [A]UMAP plot of single-cell transcriptomic profiles based on groups, cell clusters and cell types. [B]Bar plot comparing the proportions of each cell type within each sample group. [C]GO biological process results obtained by top50 marker genes in each cell type. [D]Dot plot showing expression of Tight Junction markers in each group. [E]Heatmap showing the number of interactions inferred between groups.

Cells were primarily categorized into three major compartments within the intestinal mucosa: epithelial cells, stromal cells, and immune cells (Figure S1B). Epithelial cells, comprising 73.6% of the sampled cells and predominantly expressing Epcam, play crucial roles in nutrient digestion and absorption, pathogen defense, and hormone secretion(18). The underlying stromal layer’s fibroblasts, expressing collagen markers (Col1a1 or Col11a1), provide structural and nutritional support to the epithelial cells(19). Immune cells in the gut, constituting 22.4% of the population and marked by Cd45 (Ptprc), are vital for maintaining mucosal homeostasis in response to microbial infections and physical or chemical stimuli(20). This compartment includes a variety of immune cells such as T cells, B cells, plasma cells, macrophages, and mast cells. Additionally, endothelial cells expressing Pecam1 and Cdh5 and smooth muscle cells (SMC) expressing Actg2 were identified among the stromal cells (Figure 2B and S1C). Interestingly, cluster C17, characterized by its unique expression of pancreatic enzyme-related genes (e.g., Cpa1, Cpa2, Ctrb1), appeared to be derived from pancreatic acinar cells and was only found in a single sample (Figure S1D), underscoring its non-intestinal origin. Consequently, this cluster was excluded from further analyses to ensure a more accurate and reliable understanding of intestinal cell characteristics.

Further analysis based on the top 50 markers for each cell type revealed significant functional enrichment for immune and inflammatory response pathways in immune cells, while epithelial cells showed activation of antimicrobial and defensive responses. Stromal cells, including endothelial cells involved in angiogenesis, fibroblasts related to extracellular matrix and wound healing, and smooth muscle cells associated with muscle contraction, indicated distinct functional engagements (Figure 2C). This suggests that different cell types may activate specific biological processes and signaling pathways to fulfill unique functions in response to *Chlamydia* infection.

Particular attention was paid to the expression of tight junction (TJ) related markers in epithelial cells. Post-infection, a slight decrease in the expression of certain TJ genes, notably claudins (Cldns), occludin (Ocln), and junctional proteins (Tjps), was observed (Figure 2D). This highlights a disruption in the intestinal epithelial barrier, crucial for maintaining mucosal integrity, and suggests increased permeability that may facilitate bacterial pathogen penetration and subsequent infection(21).

Moreover, the interaction among various cell populations in healthy intestinal mucosa contributes to the functionality of the gut barrier. Misinterpretation of molecular signals or improper cell interactions might lead to barrier impairment and disease progression(22). Post-infection, besides the intrinsic changes within epithelial cells, the quantity of interaction signals between epithelial and other immune cell groups also showed significant alteration (Figure 2E). Notably, the interactions involving mast cells and macrophages, crucial for immune surveillance and pathogen phagocytosis(23), were diminished. This reflects the crucial role of altered immune-epithelial interactions in maintaining intestinal homeostasis following pathogen infection.

Overall, these results provide a comprehensive panorama of the unique cellular composition and alterations in the mouse colon following *Chlamydia* infection, illustrating how various cell groups cooperatively regulate epithelial barrier homeostasis, innate immunity, tissue repair, and disease progression.

### scRNA-seq Reveals Type-Specific Transcriptional Responses in Intestinal Epithelial Cells after Chlamydial Infection

Intestinal epithelial cells serve as the primary barrier against pathogen invasion, with each cell lineage playing a distinct role in pathogen resistance(24). To further delineate this, we isolated all epithelial cells from the colon and performed a detailed clustering, identifying nine major epithelial cell types. These include absorptive enterocytes (EC), transient-amplifying cells (TA), mucus- secreting goblet cells (GC), hormone-producing enteroendocrine cells (EEC), antimicrobial peptide-secreting Paneth cells (PC), pathogen-sensing tuft cells (TC), antigen-transporting microfold-like cells (M), enterocyte progenitors (EP), and intestinal stem cells (ISC) (Figure 3A). ECs, as the predominant cell type in the colon, are further categorized into proximal (EC_Proximal) and distal cells (EC_Distal) based on region-specific gene expression, which reflects their functional diversity (Figure 3B). For instance, proximal cells are specialized in absorbing iron and nutrients, whereas distal cells predominantly absorb bile acids and vitamin B12. Notably, the colonic crypts are rich in GC and TA cells located at the base, with TA cells being rapidly proliferating progeny of stem cells that give rise to ECs or EPs, and subsequently differentiate into mature ECs, GCs, and other cell types. This proliferative and differentiative capacity is vital for maintaining the integrity and renewal of the colonic mucosal layer.

**Figure 3:**
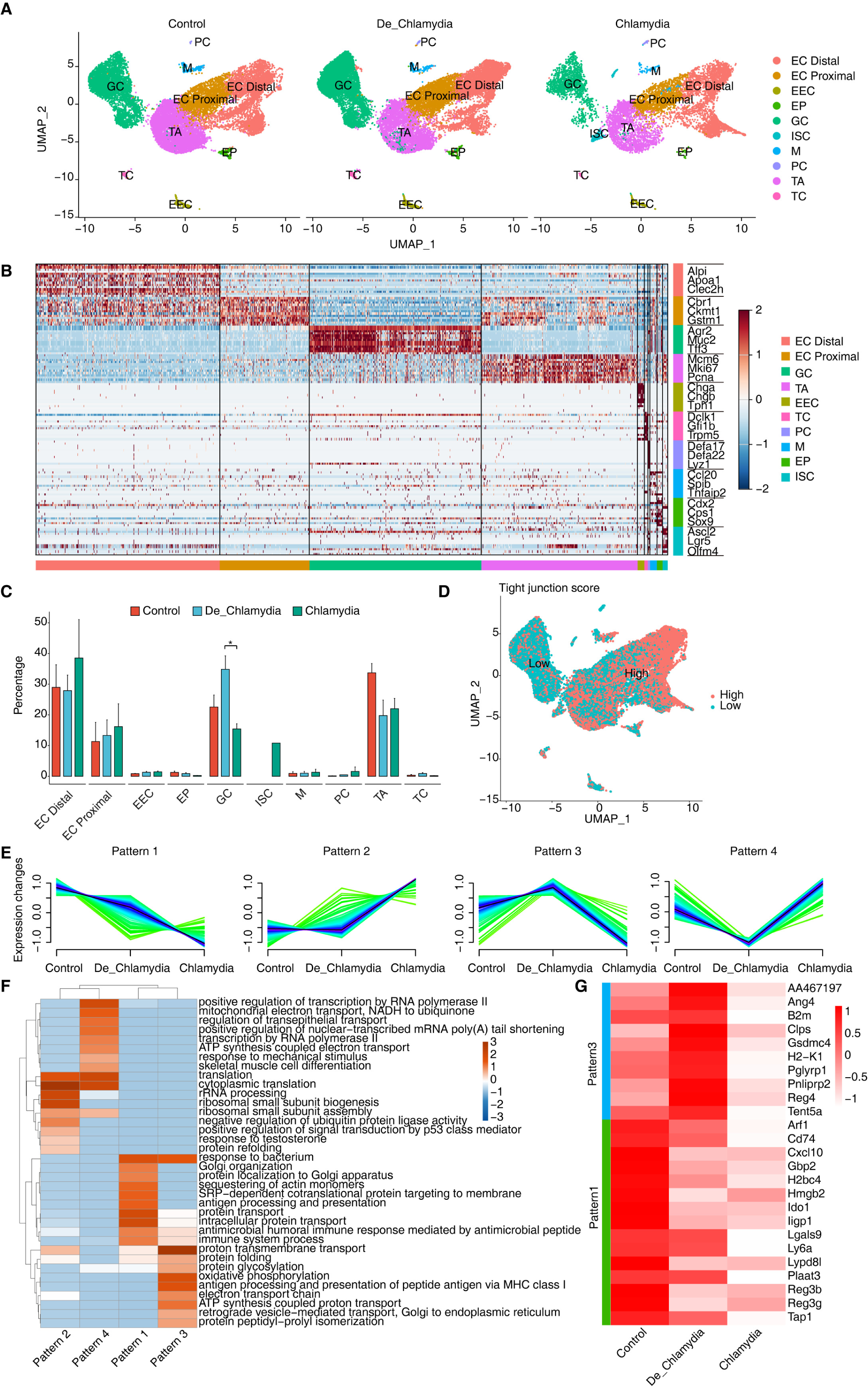
ScRNA-seq reveals intestinal epithelial cell type-specific transcriptional response evoked by *Chlamydia*l infection. [A] UMAP plots shows the composition changes of intestinal epithelial cells in different groups. [B] Heatmap shows the relative expression levels of representative markers in each cell population. [C] Bar plot compares the proportion of each cell type within each sample group. [D] UMAP plot shows the distribution score of average expression of tight junction gene set. [E] Expression patterns of differentially expressed genes in GC cells across groups base on Mfuzz. [F] GO biological process results of genes obtained from different gene expression patterns in GC cell cells. [G]Heatmap shows the expression of genes related to anti-bacterial or defense responses in different groups of GC cells.

Post-*Chlamydia* infection, a dramatic reduction in the number of TA cells was observed, suggesting a potential compromise in the epithelial renewal and repair capabilities of the gut(25). Conversely, an increase in EC numbers may represent a compensatory mechanism where, under infection or damage, ECs proliferate rapidly to replace damaged cells, thereby maintaining the integrity and functionality of the intestinal barrier(26). Importantly, we noted an increase in GC cells following heat-inactivated *Chlamydia* infection but a significant decrease in non-inactivated *Chlamydia* infections (Figure 3C). Considering the primary function of GC cells in secreting mucus and antimicrobial substances to prevent pathogen invasion and clear infected cells(27), this phenomenon suggests that while the heat-inactivated *Chlamydia* may still act as an antigenic stimulus provoking a protective response, the virulent *Chlamydia* strains might directly suppress GC proliferation.

Further analysis of TJ gene expression averages across different epithelial cell types revealed a notably lower TJ score in GC cells (Figure 3D), suggesting that *Chlamydia* might initially invade GC cells, thereby gradually compromising the integrity of the intestinal barrier. Subsequent analysis of differential gene expression within GC cells identified four expression patterns associated with *Chlamydia* infection, with patterns 1 and 3 composed of downregulated genes, and patterns 2 and 4 of upregulated genes (Figure 3E). GO enrichment analysis indicated significant enrichment of the "response to bacterium" pathway in patterns 1 and 3 (Figure 3F), implying that these downregulated genes play a proactive role in defending against *Chlamydia* infection. Focusing on antimicrobial and defense-related downregulated genes, angiogenin 4 (Ang4) emerged as the most significantly downregulated gene in the *Chlamydia* vs. Control comparison. Ang4, known for its antimicrobial properties and crucial role in maintaining intestinal epithelial homeostasis, appears to be strongly suppressed by *Chlamydia* to evade host defenses. However, its expression showed an upward trend in the De_*Chlamydia* group, reflecting a possible adaptive response of host cells to inactivated pathogens. Moreover, a consistent downregulation of antimicrobial genes such as Reg3b and Reg3g from the inactivated to the virulent *Chlamydia* infection stages indicates a weakened defensive capability of the host against pathogen invasion. Notably, the specific downregulation of the Ido1 gene in GC cells post-infection, which encodes indoleamine 2,3-dioxygenase involved in tryptophan catabolism and immune regulation(28), ranked among the top three significantly downregulated genes in both the De_*Chlamydia* and *Chlamydia* groups (Figure 3G and S2), pinpointing it as a potent target following *Chlamydia* infection in GC cells.

Overall, these findings demonstrate that *Chlamydia* infection severely impairs the functionality of intestinal epithelial GC cells, potentially by suppressing the expression of multiple genes related to antimicrobial and host defense responses, thereby facilitating the pathogen’s penetration and colonization of the epithelial barrier.

### Dysregulation of Transcriptional Regulatory Networks in Epithelial Cells During *Chlamydia* Infection

The stability of transcriptional regulatory networks is crucial for maintaining normal function in host cells. When pathogens invade, these networks can flexibly adjust the expression levels of antimicrobial or bactericidal genes to meet various environmental challenges and survival pressures, often under complex and finely tuned transcriptional control(29). To investigate potential transcriptional regulatory mechanisms in epithelial cells following *Chlamydia* infection, we utilized SCENIC to identify specific regulons, including transcription factors (TFs) and their target genes.

These regulons likely coordinate cellular physiological and pathological processes and participate in responses to *Chlamydia* infection. By calculating the Regulon Specificity Score (RSS), we found that different intestinal epithelial cell groups contain specific activated regulons, such as Brca1(+) in TA cells and Foxa3(+) in GC cells. Significant variations were also noted among different regional ECs, such as Dbp(+) in proximal cells and Ikzf2(+) in distal cells, highlighting the diversity and specificity of transcriptional regulation in intestinal epithelial cells (Figure 4A and S3A).

**Figure 4:**
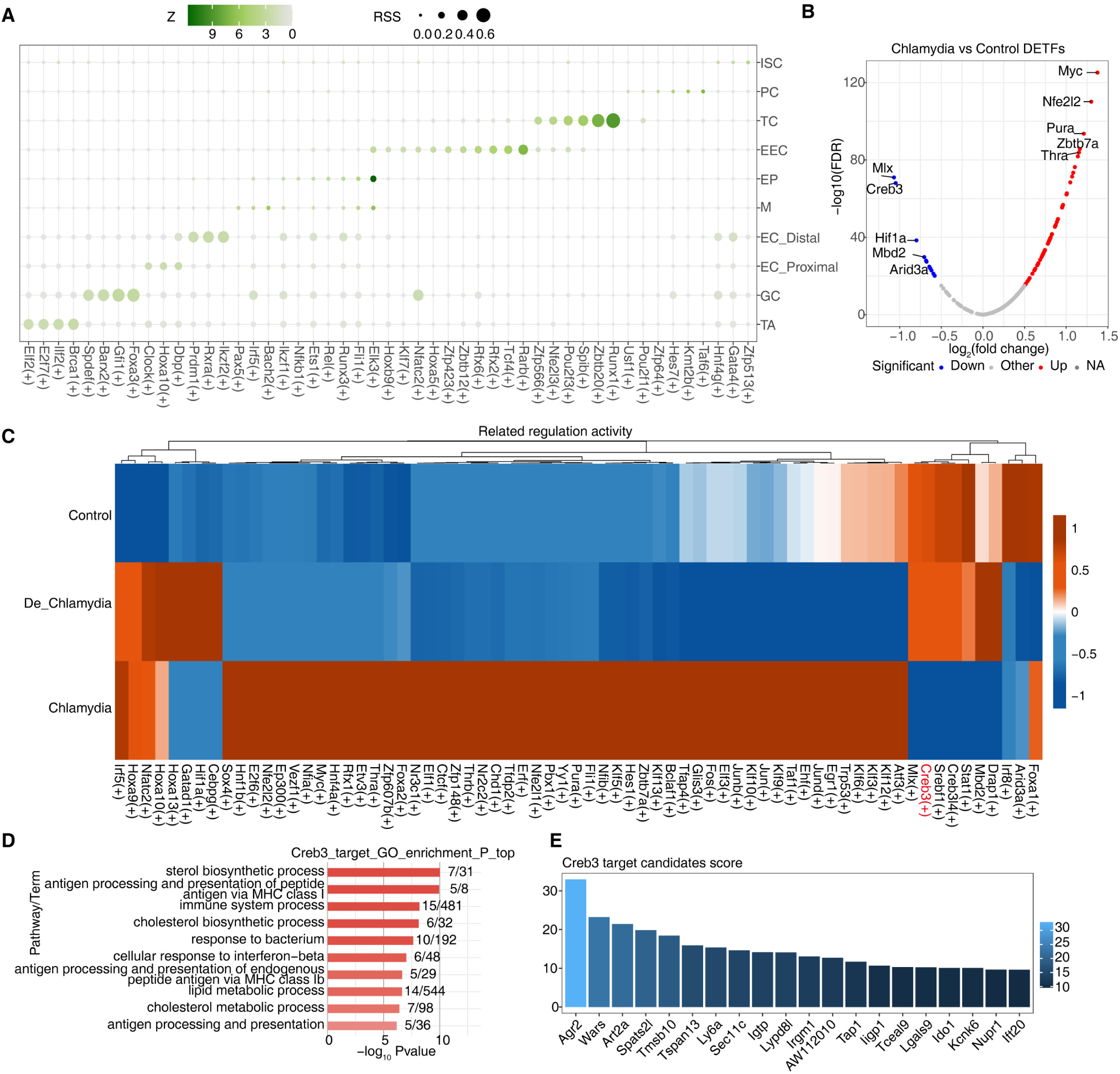
Dysregulation of transcriptional regulatory networks plays an important role in defense against *Chlamydia* infection. [A] The dot plot showing the cell-specific regulons using the Regulon Specificity Score (RSS). [B] The volcano plot showing differential transcription factors of GC cells between *Chlamydia* vs Control. Blue represents down-regulation, red represents up-regulation. [C]The heatmap showing AUC regulatory activity of *Chlamydia* vs Control DETFs in GC cells of different groups. [D]GO-enriched pathways of target DEGs regulated by Creb3 in GC cells. [E]The bar plot shows the importance scores of top 20 target genes regulated by Creb3 in GC cells.

Further analysis of their target genes through GO enrichment demonstrated their involvement in dissimilar biological functions, indicating not only their activity in specific cell types or regions but also their functional specialization (Figure S3B). We particularly focused on GC cells, which in the *Chlamydia* vs. Control comparison revealed 80 TFs with significantly differential expression—68 upregulated and 12 downregulated (Figure 3B). In contrast, only 27 differentially expressed TFs were detected in the De_*Chlamydia* vs. Control comparison, suggesting that the impact of inactivated *Chlamydia* infection on the transcriptional regulatory network stability of GC cells is less severe than that of viable *Chlamydia*.

Examination of these differentially expressed TFs in the three groups of GC cells showed that downregulated TFs had more pronounced changes in AUC activity post-infection, particularly Creb3 (Figure 4C and S3D). As a member of the cAMP response element-binding protein family, Creb3 plays a pivotal role in the unfolded protein response (UPR), a critical component of cellular defense mechanisms often induced by pathogenic invasion that affects protein folding. Disruption in Creb3 transcriptional regulatory mechanisms could be significantly associated with *Chlamydia* infection(30). To understand whether Creb3 is involved in regulating the expression of host cell genes related to anti-infection, GO analysis revealed that Creb3-regulated target genes are significantly enriched in responses to bacteria and interferon-beta (Figure 4E), suggesting Creb3’s vital regulatory role in antimicrobial and antiviral defense processes. KEGG pathway analysis further indicated that Creb3-regulated target genes are primarily associated with endoplasmic reticulum protein processing and viral infection pathways (Figure S3E), emphasizing Creb3’s critical role in regulating cellular ER stress and UPR, whose mediated transcriptional regulatory network disruption might be a key pathogenic mechanism allowing *Chlamydia* to invade GC cells.

Additionally, based on the regulatory strength scores of target genes regulated by each TF, we found that Creb3 might strongly regulate the expression of numerous antimicrobial genes in GC cells, such as Ly6a, Lypd8l, Tap1, Ligp1, Lgals9, and specifically the downregulated Ido1 (Figure 4D), further showcasing Creb3’s importance in combating *Chlamydia* infection and host defense. Moreover, we also discovered a significant positive correlation between Creb3 and Ido1 expression in GC cells (Figure S3F), which is thought to possibly constrain the specific expression of Ido1 in GC cells by inhibiting Creb3 activity post-*Chlamydia* infection, leading to impaired GC cell barrier function.

These findings indicate that *Chlamydia* infection leads to a disruption of multiple functional TF-mediated gene regulatory networks in GC cells, particularly those related to UPR and involving Creb3. By regulating differential expression of host defense and anti-infection-related target genes, Creb3 potentially impacts the stability of the intestinal barrier.

### Alteration of Cell-Cell Interactions During *Chlamydia* Infection

The integrity of intestinal epithelium plays a critical role in inflammatory diseases and intestinal infections. The maintenance of epithelial homeostasis depends not only on the coordinated actions among different epithelial cell types but also relies on the multitude of immune cells residing within the intestine(26). During pathogenic invasion, epithelial cells secrete a vast array of signaling molecules to recruit immune cells for immunoregulation, a process vital for clearing pathogens and preventing the spread of infections(31). Moreover, the crosstalk between immune and epithelial cells is crucial for regulating epithelial proliferation, differentiation, and the restoration of barrier functions, as described in Figure 2E.

We constructed interaction networks between epithelial and immune cell groups under different infection statuses (Figure 5A). We observed robust maintenance of appropriate interactions among various epithelial cell groups and between these cells and immune cells, demonstrating an efficient cooperative mode to sustain the integrity of the epithelial barrier. Compared to the control group, interactions among epithelial cell groups, especially with the proliferative TA cells, were notably enhanced in the *Chlamydia* group. This suggests that *Chlamydia* infection may indirectly stimulate rapid renewal and regeneration of epithelial cells. Conversely, the interactions between epithelial and immune cell groups, particularly with macrophages and mast cells, were significantly reduced.

**Figure 5:**
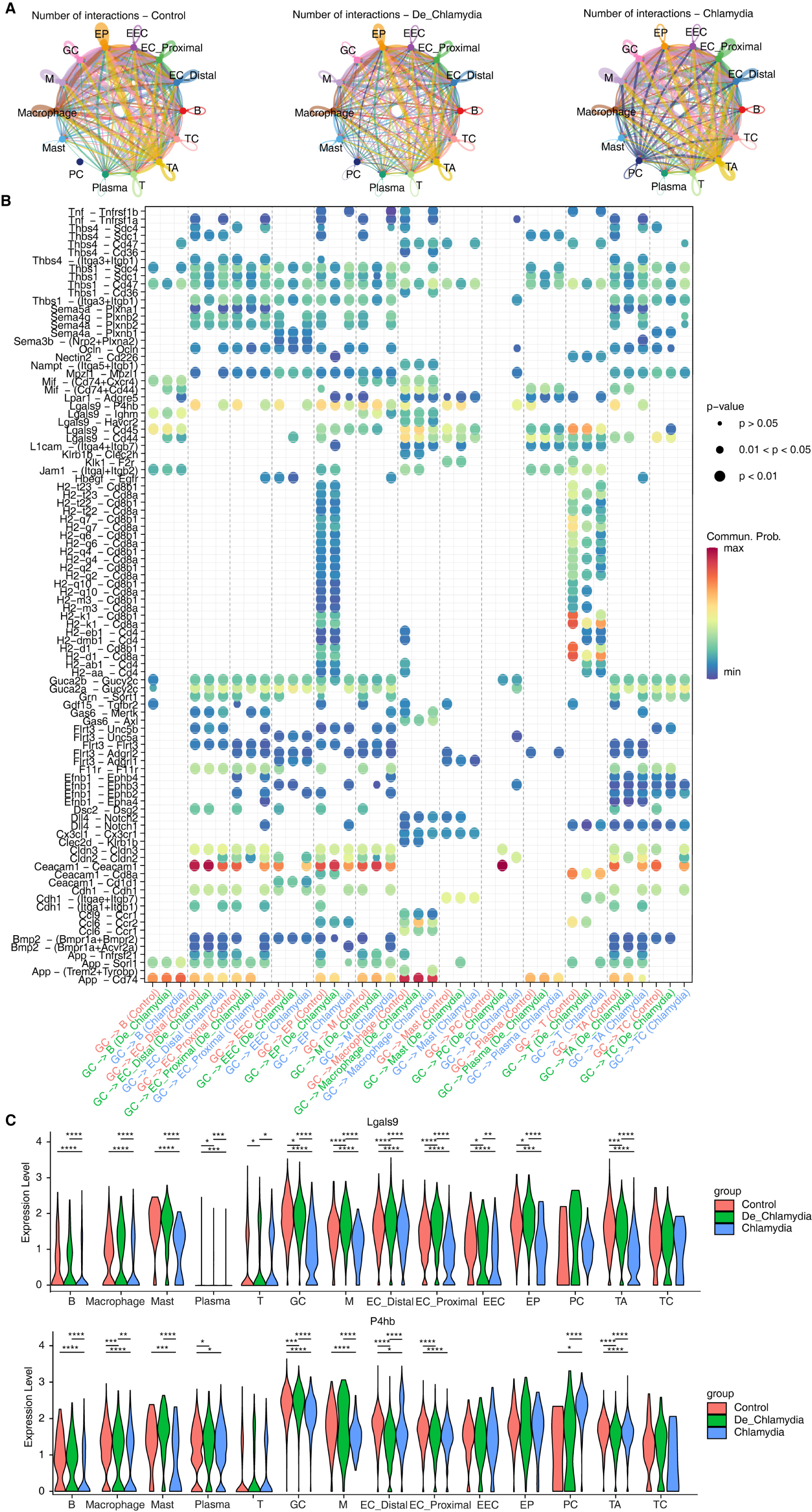
Alteration of cell-cell interactions during *Chlamydia* infection. [A]The interactions of different intestinal cell populations in each group. [B]The bubble plot shows the signaling pathways changes of GC cells with other cell types in different group. [C]The violin plot shows the expression difference of Lgals9 and P4hb in different cell types of each group. * P value ≤ 0.05, ** P value ≤ 0.01, *** P value ≤ 0.001, **** P value ≤ 0.0001.

Macrophages, known for their potent phagocytic abilities to ingest and clear pathogens including *Chlamydia*, and mast cells, which continuously monitor for pathogen invasion and attract additional immune cells to respond, illustrate that *Chlamydia* infection likely impedes this cellular communication(32), making it challenging for the host to recruit macrophages or mast cells to the infection sites for pathogen clearance.

Furthermore, we specifically highlighted the interaction patterns of goblet cells (GC) during *Chlamydia* infection, emphasizing the differences in information flow between GC cells and other types of epithelial or immune cells (Figure 5B and S4A). Post-infection, the Ceacam1-Ceacam1 signaling between GC cells and other epithelial cells was significantly weakened. Similarly, signals related to TJs (such as Ocln, Cldns) also exhibited a similar weakening, potentially indicating a substantial reduction in adhesive capacity among epithelial cells, diminishing intestinal barrier function (Figure 5B and S4B-C). In the signaling pathways involving interactions with immune cells, nearly all ligand-receptor pairs were suppressed after *Chlamydia* infection, potentially preventing GC cells from recruiting immune cells to clear the pathogen. Notably, we observed significant suppression in the Lgals9-P4hb signaling pathway within interactions involving macrophages and mast cells (Figure 5B). Lgals9 plays a critical role in resisting microbial invasions and viral infections; its significant downregulation suggests a weakened intestinal defense against *Chlamydia* infection. Importantly, the substantial downregulation of the receptor P4hb, which encodes a protein involved in preventing endoplasmic reticulum stress (ERS) and incorrect protein folding, was notable. Correlation analysis further confirmed a strong positive relationship between P4hb expression and Creb3 (Figure S4D), suggesting that P4hb might activate Creb3-mediated transcriptional regulatory networks associated with anti-infection, therefore, the Lgals9-P4hb signaling pathway in intestinal GC might have substantial potential against *Chlamydia* infection and immune evasion.

These results reveal that *Chlamydia* infection in intestinal epithelium leads to dysregulation in transcriptional regulatory homeostasis related to pathogen defense in GC cells while simultaneously suppressing the recruitment capabilities of GC cells, particularly those involving macrophages and mast cells. This maintenance likely relies on the signaling mediated by Lgals9-P4hb. *Chlamydia* facilitates long-term colonization and persistent infection in host cells by inducing the inactivation of Creb3-mediated transcriptional control related to ERS and UPR in GC cells, thereby downregulating various genes associated with pathogen defense or host immune responses, such as Ido1. This collaborative action ultimately allows *Chlamydia* to establish long-term colonization within the host cells.

### Effects of *Chlamydia* Infection on Intestinal Epithelium Cell Fate

High rates of cell turnover and plasticity contribute to the intestine’s natural adaptability, maintaining homeostasis and health through the differentiation of various epithelial cell lineages(33). However, the impact of *Chlamydia* infection on epithelial lineage differentiation and cell fate remains unclear. In this study, we delineated the lineage structure of intestinal epithelial cells and their pseudo-time changes to simulate the differentiation process. Initially, we inferred four potential fate trajectories (lineages 1-4) from multipotent progenitor cells (EP) to fully differentiated cells. Lineages 1-3 respectively showed the differentiation of TA cells into goblet cells (GC), tuft cells (TC), and enteroendocrine cells (EEC). Lineage 4 likely represents the cell fate of enterocytes (EC) (Figure 6A). As anticipated, post-*Chlamydia* infection, a noticeable reduction in TA and GC cell numbers was observed along with a pronounced slowing of differentiation in lineage 1 as pseudo-time progressed (Figure 6B-C).

**Figure 6:**
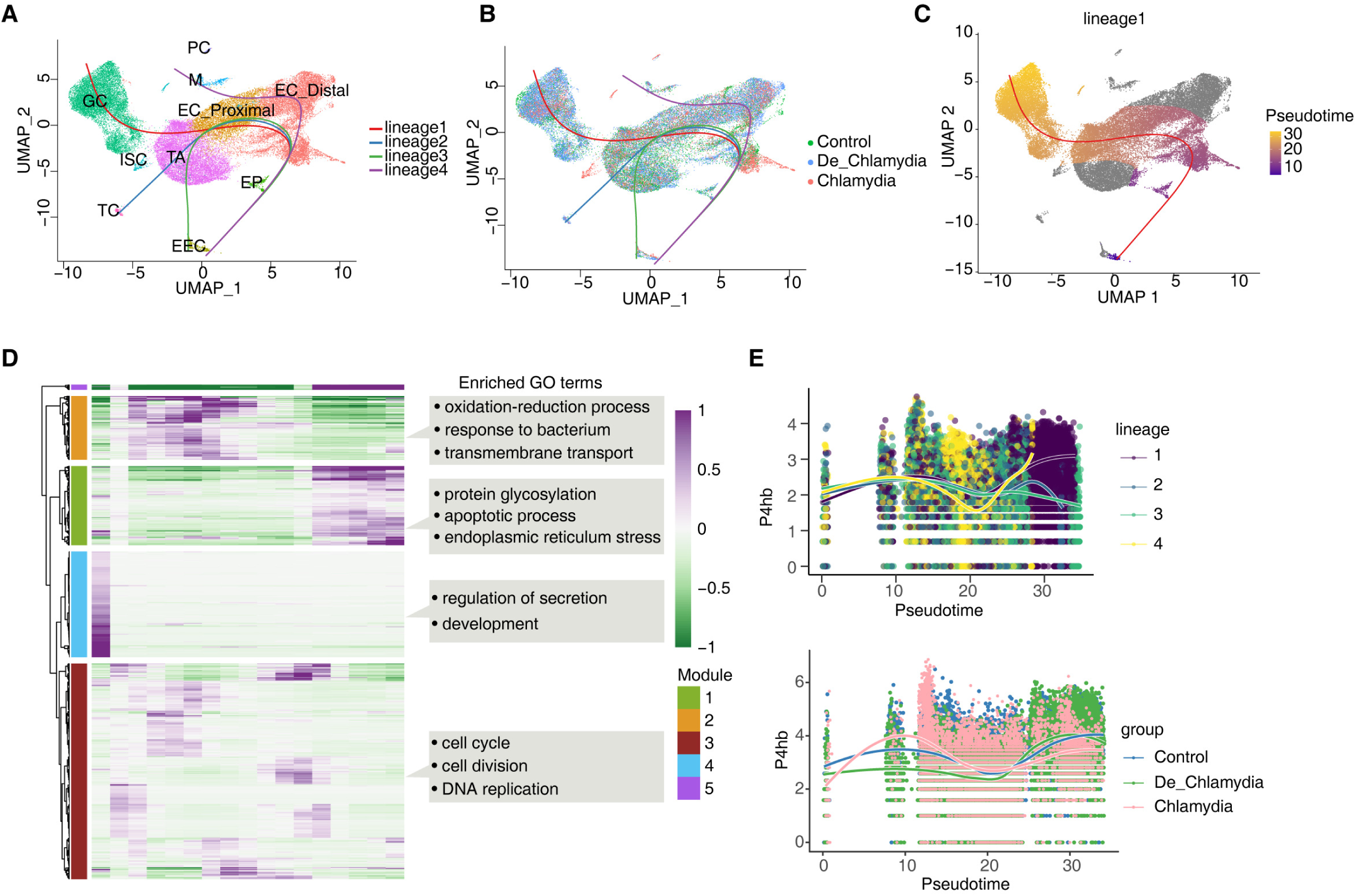
Effects of *Chlamydia* infection on intestinal epithelium cell fate. [A] UMAP plot shows the differentiation trajectories of intestinal epithelial cell lineages.[B] UMAP plot shows the distribution of differentiation trajectories in different groups. [C] UMAP plot shows the differentiation trajectory of GC (lineage 1). [D]Heatmap shows pseudotime expression changes and clustering modules of genes in lineage 1. [E]The pseudotime expression changes of P4hb in different lineages and its expression differences among different groups in lineage 1.

We further examined gene expression changes along lineage 1’s trajectory, clustering them into four gene expression modules (Modules 1-4), where Modules 1 and 5 genes exhibited strong expression in the later stages of pseudo-time, crucial for determining the final fate of GC cells. GO analysis revealed that Module 1 genes were enriched in processes related to protein glycosylation, apoptosis, and endoplasmic reticulum stress (ERS) (Figure 6D and S5A), highlighting ERS’s significant role in mediating the differentiation process of the GC lineage. Module 5 genes predominantly represented GC-specific genes, such as the antimicrobial peptide Ang4 and the mucin Muc2, which together form a robust mucus barrier.

Finally, we focused on the pseudo-time expression changes of P4hb and Creb3, which are involved in ERS within GC. Results showed that P4hb exhibited strong upregulation in the later stages of the GC lineage, closely associated with GC cell fate. However, in the *Chlamydia* group, P4hb showed significantly lower expression, suggesting that *Chlamydia* infection might suppress P4hb, significantly inhibiting GC cell lineage differentiation and leading to a reduction in the number of intestinal GC cells (Figure 6D). In contrast, the pseudo-time changes of Creb3 were relatively minor (Figure S5B), likely more related to the homeostasis and function of fully differentiated GC cells.

In summary, *Chlamydia* infection might suppress the expression of P4hb, thereby weakening GC cell proliferation and reducing the recruitment efficiency of immune cells. This process further hinders the activation of key anti-infection transcriptional regulatory networks mediated by Creb3 within GC cells, limiting the expression of anti-infection related genes. The significant reduction in GC cell-secreted antimicrobial proteins ultimately provides an opportunity for *Chlamydia* to breach the intestinal epithelial barrier, successfully establish colonization, and trigger a range of host infection symptoms.

## DISCUSSION

*Chlamydia trachomatis* (*C. trachomatis*) has long been recognized as a leading sexually transmitted infection with severe consequences for female reproductive health(1). Recent evidence suggests that beyond the genital tract, the GI tract serves as a significant reservoir for the persistence of *Chlamydia*(5). *Chlamydia* is routinely detected in the human GI tract according to clinical studies(3). *C. trachomatis* can spread to the GI tract through sexually independent pathways, regardless of oral or anal sex history(4). Our study builds on the growing understanding that *Chlamydia* can establish long-term colonization in the large intestine, which may serve critical roles in both the pathogenesis of the bacteria and the chronicity of the infection(5).

*Chlamydia* is believed to inhabit the gut through mechanisms that involve the evasion of host immune responses and the exploitation of plasmid-related resistance against the killing mechanisms of the GI tract(7). Research suggests that the process of *Chlamydia* spreading to and colonizing the GI tract occurs by steps. Certain mutants of *Chlamydia muridarum* display reduced capabilities in colonizing the GI tract; for instance, mutants lacking plasmid functions exhibit significant difficulties in colonizing the upper GI tract, whereas those deficient in specific chromosomal genes show more pronounced obstacles in the lower GI tract(34). This observation indicates that *Chlamydia* may depend on plasmids for its dissemination to the large intestine, while chromosome-encoded factors are necessary for sustaining colonization(7). Despite the presence of various pathogenic virulence factors of *Chlamydia*, additional research is warranted to further investigate the mechanism of how *Chlamydia* colonizes the host within the gut.

Our investigation confirms the large intestine as the primary site for *Chlamydia* colonization, aligning with observations that *Chlamydia* detections in the rectal swabs of mammals(35). This localization is particularly important in understanding the pathogen’s tissue preference and its ability to evade standard detections that focus mostly on the genital tract(36). The persistence of *Chlamydia* in the gut necessitates a broader approach in treatment strategies that target this additional reservoir to effectively manage or eradicate the infection.

The dynamic alterations in the cellular composition of the mouse colon long term infection, as observed in our results, underscore the infection’s impact on gut homeostasis. Significant changes in the proportion and function of epithelial cells, including goblet and TA cells, highlight the infection’s capability to disrupt normal cellular turnover and barrier functions(16) . These changes are critical as they directly compromise the intestinal barrier, facilitating easier pathogen penetration and wider spread within the host. Particularly, the functionality of goblet cells, which are essential in forming the mucosal barrier against pathogens(27), is severely compromised under *Chlamydia* infection. The suppression of key antimicrobial and host defense genes in these cells not only facilitates the pathogen’s colonization but also indicates potential targets for therapeutic intervention to reinforce the barrier and prevent infection progression.

The stability of transcriptional regulatory networks within these cells plays a pivotal role in their ability to respond to environmental stresses and pathogenic challenges(29). Our findings that *Chlamydia* disrupts these networks, particularly through the alteration of key transcription factors like Creb3, suggest that the pathogen employs sophisticated mechanisms to dampen the host’s immune response(37). This disruption likely provides a conducive environment for the pathogen to thrive, underscoring the complexity of *Chlamydia*’s interaction with host cellular machinery.

Furthermore, our results showed the possible crucial role of altered cell-cell communication in degrading intestinal barrier integrity and modulating immune responses. The weakening of specific signaling pathways, such as the Lgals9-P4hb pathway in goblet cells, marks a significant mechanism by which *Chlamydia* evades host defenses. This finding suggests that bolstering such pathways could enhance the gut’s defensive response and potentially curb the spread of infection.

Finally, the impact of *Chlamydia* on the differentiation trajectories of intestinal epithelial cells, particularly affecting cell fate and lineage commitment, highlights another layer of complexity in the pathogen-host interaction. The suppression of P4hb expression, crucial for the proliferation and function of goblet cells(38), suggests that *Chlamydia* strategically impairs cellular defenses and regeneration capabilities to facilitate its colonization and persistence.

Despite the significant insights provided by this study, there are several limitations that warrant consideration. Firstly, while scRNA-seq was employed to detect and analyze gene expression changes associated with *Chlamydia* infection, this method alone cannot fully establish the functional roles of these changes. No additional validation experiments, such as genetic manipulation or pharmacological intervention, were conducted to confirm the findings directly. This limits our ability to ascertain the causal relationships between gene expression changes and observed phenotypes.

Additionally, this study exclusively utilized a mouse model to investigate the colonization and impact of *Chlamydia* in the GI tract. While mouse models are invaluable for understanding disease mechanisms due to their physiological similarities to humans and their ease of genetic manipulation, the results may not fully translate to human biology. The complex interactions between *Chlamydia* and its host that were observed in mice require validation in human tissues or through clinical studies to ensure their relevance to human health. Further validation through dedicated animal studies and eventual confirmation in human subjects will be crucial to solidify these biomarkers and confirm their biological relevance across different species.

In summary, the results presented in this study provide a detailed panorama of the cellular and molecular alterations in the colon following *Chlamydia* long term infection, illustrating the intricate mechanisms by which the pathogen manipulates host cellular functions to establish a secure niche within the GI tract. These insights not only contribute to our understanding of *Chlamydia* as a multifaceted pathogen but also underscore the necessity for integrated approaches to tackle this infection, considering its ability to colonize sites beyond the primary site of infection. The potential for targeted interventions aimed at restoring gut integrity and enhancing host resistance offers promising avenues for research and therapeutic development.

## MATERIALS AND METHODS

### Animals

C57BL/6J female mice, aged 5-6 weeks, were sourced from Vital River in Beijing and maintained under controlled environmental conditions (temperature: 22 °C; light/dark cycle: 12 hours each). Following a one-week acclimatization period in the laboratory, these mice were used for experiments that received approval from the Ethics Committee of the Institute of the 3^rd^ Xiangya Hospital, Central South University, adhering to the guidelines set by the Chinese Council on Animal Care.

### Preparation of *Chlamydial* Organisms

The *C. muridarum* strains used in this research were derived from the Nigg3 strain (GenBank accession number CP009760.1). We cultured *Chlamydia* in HeLa cells, followed by purification of elementary bodies (EBs) using established protocols. These EBs were aliquoted and stored at -80 °C. For deactivation, EBs underwent a 30-minute heat treatment at 56 °C.

### Inoculation and Monitoring of *Chlamydia* Infections in tissues and swabs

Mice aged 6 to 7 weeks received intragastric inoculations of purified *C. muridarum* EBs (either live or heat deactivated), with each mouse receiving 2 × 10^5 inclusion-forming units (IFUs). The naïve control group was administered a sucrose-phosphate-glutamic acid (SPG) vehicle.

To monitor the shedding of live organisms, anorectal swabs were collected different time points after inoculation. Each swab was immersed in 0.5 ml of SPG and vortexed with glass beads to extract *Chlamydial* organisms, which were then titrated on HeLa cell monolayers in duplicate. The infected cultures were processed for immunofluorescence assays as described below. Inclusions were counted in five random fields per coverslip under a fluorescence microscope. For coverslips with fewer than one IFU per field, the entire coverslip was counted. Coverslips exhibiting significant cytotoxicity in HeLa cells were excluded from analysis. The total number of IFUs per swab was calculated based on the average IFUs per view, the area ratio of the view to that of the well, dilution factors, and inoculation volumes. If applicable, mean IFUs per swab were derived from serially diluted and duplicate samples. The total count of IFUs per swab was converted to log10 and used to compute the mean and standard deviation for the mice within each group at each time point, as previously published(39).

For quantitating live organisms from mouse GI segments, each organ or tissue segment was transferred to a tube containing 0.5 to 5 ml SPG depending the sizes of the organs(39). GI tract tissues include esophagus, stomach, small intestine (duodenum, jejunum, and ileum), and large intestine (cecum, colon, and rectum). The tissue segments were homogenized in cold SPG using a 2-ml tissue grinder or an automatic homogenizer. The homogenates were briefly sonicated and spun at 3,000 rpm for 5 min to pellet remaining large debris. After sonication, the supernatants were titrated for live *C. muridarum* organisms on HeLa cells as described above. The results were expressed as log10 IFU per tissue segment.

### Quality control of scRNA-seq data

The scRNA-seq data included 59,197 cells from 3 Control samples, 3 De_*Chlamydia*-treated samples and 3 *Chlamydia*-treated mouse large intestine tissue samples. The UMI count matrix was converted into a Seurat object by the R package Seurat (version 4.3.0)(40). Cells with UMI numbers <1000 or with detected genes < 500 or with over 20% mitochondrial-derived UMI counts were considered low-quality cells and were removed. Genes detected in less than 5 cells were removed for downstream analyses.

### scRNA-seq data preprocessing

After quality control, the UMI count matrix was log normalized. Then top 2000 variable genes were used to create potential Anchors with FindIntegrationAnchors function of Seurat. Subsequently, IntegrateData function was used to integrate data. To reduce the dimensionality of the scRNA-Seq dataset, principal component analysis (PCA) was performed on an integrated data matrix. With Elbowplot function of Seurat, top 50 PCs were used to perform the downstream analysis. The main cell clusters were identified with the FindClusters function offered by Seurat, with resolution set as default (res = 0.4). Finally, cells were clustered into 10 major cell types. And then they were visualized with UMAP plots. To identify the cell type for each cluster, we detected gene markers for each cell clusters using the “FindAllMarkers” function in Seurat package (v4.3.0) on a natural log scale was at least 0.5 and the difference of percent of detected cells was at least 0.25 and adjusted p-value was less than 0.05, then we annotated cell types using ScType tools(41).

### Differential gene expression analysis

Differentially expressed genes (DEGs) were determined with the FindMarkers / FindAllMarkers function from the Seurat package (one-tailed Wilcoxon rank sum test, pvalues adjusted for multiple testing using the Bonferroni correction). For computing DEGs, all genes were probed that the expression difference on a natural log scale was at least 0.5 and the difference of percent of detected cells was at least 0.15 and adjusted p-value was less than 0.05.

### Transcription factor regulatory network analysis

The modules of TFs were identified by the SCENIC (42) python workflow (version 0.11.2) using default parameters (http://scenic. aertslab.org). A human/mouse TF gene list was used from the resources of pySCENIC (https://github.com/aertslab/pySCENIC/tree/master/resources). Activated TFs were identified in the AUC matrix, and differentially activated TFs were selected using R package limma(43) based on the fold change (logFC ≥ 0.5 or ≤ -0.5) and false discovery rate (FDR ≤ 0.05). To identify cluster-specific regulons (especially for analyses with many cell types, where some regulons are common to multiple of them) we used the Regulon Specificity Score (RSS)(44).

### Cell–cell communication

Cell–cell interactions based on the expression of known ligand–receptor pairs in different cell types were inferred using CellChat(45) (v2.1.1). To identify potential cell–cell communication networks perturbed or induced in mice large intestine, we followed the official workflow and loaded the normalized counts into CellChat and applied the preprocessing functions identifyOverExpressedGenes, identifyOverExpressedInteractions and projectData with standard parameters set. As database, we selected the Secreted signaling pathways and used the precompiled protein–protein-interactions as a priori network information. For the main analyses the core functions computeCommunProb, computeCommunProbPathway and aggregateNet were applied using standard parameters and fixed randomization seeds. Finally, to determine the senders and receivers in the network, the function netAnalysis_signallingRole was applied on the netP data slot.

### Functional enrichment analysis

To sort out functional categories of genes, Gene Ontology (GO) terms and KEGG pathways were identified using KOBAS 2.0(46). Hypergeometric test and Benjamini-Hochberg FDR controlling procedure were used to define the enrichment of each term.

### Pseudotime trajectory analysis

We used Slingshot(47) to infer developmental differentiation trajectories in the scRNA-seq dataset. After dimensionality reduction and clustering, Slingshot can serve as a component in an analysis pipeline by identifying the global lineage structure with a cluster-based minimum spanning tree, and fitting simultaneous principal curves to describe each lineage, and inferring pseudo-time variables.

### Gene expression pattern analysis

We performed time-dependent trend analysis of gene expression using the Mfuzz package of R language(48). First, the average expression level log2 (TPM/10 + 1) of each gene in each stage among single cells was calculated. Then, the ‘timeclust’ function was used to cluster different expression patterns.

### Other statistical analysis

Tight junction scores were calculated using the Seurat function AddModuleScore, which analyses the average expression per cell. For statistical significance testing of two independent groups, an unpaired two-tailed Student’s t-test was used. P-values of < 0.05 were considered statistically significant.

**Figure S1. ScRNA-seq reveals *Chlamydia* infection associated shift in intestinal cell composition.** [A] Dot plot showing expression of representative markers in each cell type. [B] UMAP plot shows the distribution of different cell types. [C] Pie chart shows the cell number and proportion of each cell type. [D] Histogram shows the proportion of cell populations of each cell type in each sample.

**Figure S2. ScRNA-seq reveals intestinal epithelial cell type-specific transcriptional response evoked by *Chlamydia*l infection.** [A] Volcano plot shows DEGs results of different epithelial cell populations between *Chlamydia* vs Control groups. [B] Volcano plot shows DEGs results of different epithelial cell populations between De_*Chlamydia* vs Control groups.

**Figure S3.** Dysregulation of transcriptional regulatory networks plays an important role in defense against *Chlamydia* infection. [A] UMAP plot shows the AUC activity distribution of selected regulons in epithelial cells. [B] GO biological process results of target genes regulated by TFs specific to different epithelial cells. [C] Bar graphs show the number of differentially expressed TFs compared between different groups. [D] UMAP plot shows the AUC activity of Creb3(+) in each group. [E] The most enriched KEGG pathways of target DEGs regulated by Creb3 in GC. [F] Correlation scatter plot shows the interplay between Creb3 and Ido1 expression in GC.

**Figure S4. Alteration of cell-cell interactions during *Chlamydia* infection.** [A] Interactions of GC cells in different groups. [B] CEACAM signal pathway network in different groups. [C] OCLN signal pathway network in different groups. [D] Correlation scatter plot shows the interplay between Creb3 and P4hb expression in GC.

**Fig.S5 Effects of *Chlamydia* infection on intestinal epithelium cell fate.** [A] GO enrichment results of different module genes. [B] The pseudotime expression changes of Creb3 in different lineages and its expression differences among different groups in lineage 1.

## Conflict of Interest

The authors declare that the research was conducted in the absence of any commercial or financial relationships that could be construed as a potential conflict of interest.

## Author Contributions

YCJ, SX, and CQS wrote the original draft. YYH, JW, QZ, ZZZ, and XS reviewed and edited the manuscript. LYW and TYZ contributed to the figure preparations. QT, TYZ and LYW provided the conceptualization and funding. All authors contributed to the article and approved the submitted version.

## Funding

This research is supported by National Natural Science Foundation of China for the Youth (32100162) to QT and (32000138) to TZ. The Natural Science Foundation of Hunan Province, China (2022JJ70092 and 2023JJ40355), the Science and Technology Innovation Program of Hunan Province (2024RC3233 and 2021SK4021).

## Acknowledgments

We express our gratitude to Prof. Wu Xiang from the Department of Parasitology at Central South University’s Xiangya Medical School for her support in the preparation of *Chlamydia*. Additionally, we extend our thanks to Ruixing Biotechnology Co., Ltd (Wuhan) for the scRNA-seq service, and the subsequent bioinformatics analysis, which were crucial for our research.

## Data Availability Statement

The original contributions presented in the study are included in the article/supplementary material, further inquiries can be directed to the corresponding authors.

